# A resource-rational theory of set size effects in visual working memory

**DOI:** 10.1101/151365

**Authors:** Ronald van den Berg, Wei Ji Ma

## Abstract

Encoding precision in visual working memory decreases with the number of encoded items. Here, we propose a normative theory for such set size effects: the brain minimizes a weighted sum of an error-based behavioral cost and a neural encoding cost. We construct a model from this theory and find that it predicts set size effects. Notably, these effects are mediated by probing probability, which aligns with previous empirical findings. The model accounts well for effects of both set size and probing probability on encoding precision in nine delayed-estimation experiments. Moreover, we find support for the prediction that the total amount of invested resource can vary non-monotonically with set size. Finally, we show that it is sometimes optimal to encode only a subset or even none of the relevant items in a task. Our findings raise the possibility that cognitive “limitations” arise from rational cost minimization rather than from constraints.

## INTRODUCTION

A well-established property of visual working memory (VWM) is that the precision with which items are encoded decreases with the number of encoded items (Ma et al. 2014; Luck & Vogel 2013). A common way to explain this set size effect has been to assume that there is a fixed amount of resource available for encoding: the more items, the less resource per item and, therefore, the lower the precision per item. Different forms have been proposed for this encoding resource, such as samples (Palmer 1994; Sewell et al. 2014), Fisher information (Van Den Berg et al. 2012; Keshvari et al. 2013), and neural firing rate (Bays 2014). Unless additional assumptions are made, models with a fixed amount of resource generally predict that the encoding precision per item (defined as inverse variance of the encoding error) is inversely proportional to set size. It has turned out that this prediction is often inconsistent with empirical data, which is the reason that more recent studies instead use a power law to describe set size effects (Bays et al. 2009; Bays & Husain 2008; Van Den Berg et al. 2012; van den Berg et al. 2014; Devkar & Wright 2015; Elmore et al. 2011; Mazyar et al. 2012; Wilken & Ma 2004; Donkin et al. 2016; Keshvari et al. 2013). In the more flexible power-law models, the total amount of resource across all items is no longer fixed, but instead decreases or increases monotonically with set size. These models tend to provide excellent fits to experimental data, but they have been criticized for lacking a principled motivation (Oberauer et al. 2016; Oberauer & Lin 2017): they accurately describe how memory precision depends on set size, but not why these effects are best described by a power law – or why they exist at all. In the present study, we seek a *normative* answer to these fundamental questions.

While previous studies have used normative theories to account for certain aspects of VWM, none of them has accounted for set size effects in a principled way. Examples include our own previous work on change detection (Keshvari et al. 2012; Keshvari et al. 2013), change localization (Van Den Berg et al. 2012), and visual search (Mazyar et al. 2012). In those studies, we modelled the decision stage using optimal-observer theory, but assumed an ad hoc power law to model the relation between encoding precision and set size. Another example is the work by Sims and colleagues, who developed a normative framework in which working memory is conceptualized as an optimally performing information channel (Sims 2016; Sims et al. 2012). Their information-theoretic framework offers parsimonious explanations for the relation between stimulus variability and encoding precision (Sims et al. 2012) and the non-Gaussian shape of encoding noise (Sims 2015). However, it does not offer a normative explanation of set size effects. In their early work (Sims et al. 2012) they accounted for these effects by assuming that total information capacity is fixed, which is similar to other fixed-resource models and predicts an inverse proportionality between encoding precision and set size. In their later work (Orhan et al. 2014; Sims 2016), they add to this the assumption that there is an inefficiency in distributing capacity across items and fit capacity as a free parameter at each set size. Neither of these assumptions has a normative motivation. Finally, Nassar and colleagues have proposed a normative model in which a strategic trade-off is made between the number of encoded items and their precision: when two items are very similar, they are encoded as a single item, such that there is more resource available per encoded item (Nassar et al. 2018). They showed that this kind of “chunking” is rational from an information-theoretical perspective, because it minimizes the observer’s expected estimation error. However, just as in much of the work discussed above, this theory assumes a fixed resource budget for item encoding, which is not necessarily optimal when resource usage is costly.

The approach that we take here aligns with the recent proposal that cognitive systems are “resource-rational”, i.e., trade off the cost of utilizing resources against expected task performance (Griffiths et al. 2015). The starting point of our theory is the principle that neural coding is costly (Attwell & Laughlin 2001; Lennie 2003; Sterling & Laughlin 2015), which may have pressured the brain to trade off the behavioral benefits of high precision against the cost of the resource invested in stimulus encoding (Pestilli & Carrasco 2005; Lennie 2003; Ma & Huang 2009; Christie & Schrater 2015). We hypothesize that set size effects – and limitations in VWM in general – may be the result of making this trade-off near-optimally. We formalize this hypothesis in a general model that can be applied to a broad range of tasks, analyze the theoretical predictions of this model, and fit it to data from nine previous delayed-estimation experiments.

## THEORY

### General theoretical framework: trade-off between behavioral and neural cost

We define a vector **Q**={*Q*_1_,…, *Q_N_*} that specifies the amount of resource with which each of *N* task-relevant items is encoded. We postulate that **Q** affects two types of cost: an expected behavioral cost 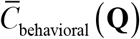 induced by task errors and an expected neural cost 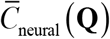 induced by spending neural resources on encoding. The *expected total cost* is a weighted combination,

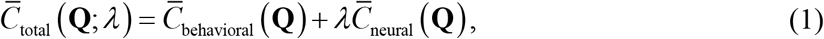

where the weight λ≥0 represents the importance of the neural cost relative to the behavioral cost. Generally, increasing the amount of resource spent on encoding will reduce the expected behavioral cost, but simultaneously increase the expected neural cost.

The key novelty of our theory is that instead of assuming that there is a fixed resource budget for stimulus encoding (a hard constraint), we postulate that the brain – possibly on a trial-by-trial basis – chooses its resource vector **Q** in a manner that minimizes the expected total cost. We denote the vector that yields this minimum by **Q**_optimal_:

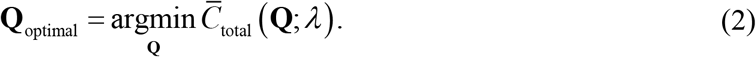

Under this policy, the total amount of invested resource – the sum of the elements of **Q**_optimal_ – does not need to be fixed: when it is “worth it” (i.e., when investing more resource reduces the expected behavioral cost more than it increases the expected neural cost), more resource may be invested.

Eqs. (1) and (2) specify the theory at the most general level. To derive testable predictions from this framework, we next propose specific formalizations of resource and of the two expected cost functions.

### Formalization of resource

As in our previous work (Keshvari et al. 2012; Keshvari et al. 2013; Mazyar et al. 2012; Van Den Berg et al. 2012; van den Berg et al. 2014), we quantify encoding precision as Fisher information, *J*. This measure provides a lower bound on the variance of any unbiased estimator (Cover & Thomas 2005; Ly, A. et al. 2015) and is a common tool in the study of theoretical limits on stimulus coding and discrimination (Abbott & Dayan 1999). Moreover, we assume that there is item-to-item and trial-to-trial variation in precision (Fougnie et al. 2012; Van Den Berg et al. 2012; van den Berg et al. 2014; Keshvari et al. 2013; van den Berg et al. 2017). Following previous work (e.g., (Van Den Berg et al. 2012; van den Berg et al. 2014)), we model this variability using a gamma distribution with a mean 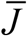 and shape parameter *τ*≥0 (larger *τ* means more variability); we denote this distribution by Gamma 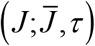.

We specify resource vector **Q** as the vector with mean encoding precisions, 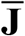, such that the general theory specified by Eqs. (1) and (2) modifies to

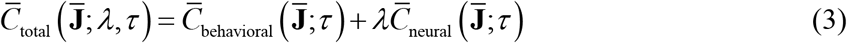

and

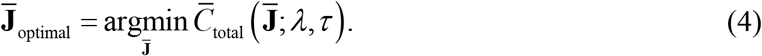

In this formulation, it is assumed that the brain has control over resource vector 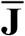, but not over the variability in how much resource is actually assigned to an item. However, our choice to incorporate variability in *J* is empirically motivated and not central to the theory: parameter *τ* mainly affects the kurtosis of the predicted estimation error distributions, not the variance of these distributions or the way that the variance depends on set size (which is the focus of this paper). We will show that the theory also predicts set size effects when there is no variability in *J*.

### Formalization of expected neural cost

To formalize the neural cost function, we make two general assumptions. First, we assume that the expected total neural cost is the sum of the expected neural costs associated with the *N* individual items. Second, we assume that each of these “local” neural costs has the same functional dependence on the amount of allocated resource: if two items are encoded with the same amount of resource, they induce equal amounts of neural cost. Combining these assumptions, the expected neural cost induced by encoding *N* items with resource 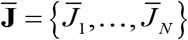 takes the form

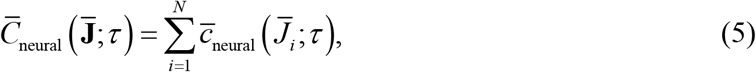

where we introduced the convention to denote local costs (associated with a single item) with small *c*, to distinguish them from the global costs (associated with the entire set of encoded items), which we denote with capital *C*.

The expected local neural cost induced by encoding an item with resource 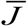 is obtained by integrating the amount of local neural cost induced by investing an amount of resource *J*, which we will denote by *c_neural_*(*J*), over *J*,

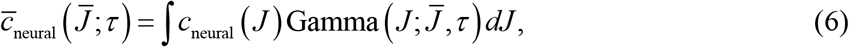

The theory is agnostic about the exact nature of the cost function *c*_neural_(*J*): it could include spiking and non-spiking components (Lennie 2003), be associated with activity in both sensory and non-sensory areas, and include other types of cost that are linked to “mental effort” in general (Shenhav et al. 2017).

To motivate a specific form of this function, we consider the case that the neural cost is incurred by spiking activity. For many choices of spike variability, including the common one of Poisson-like variability (Ma et al. 2006), Fisher information *J* of a stimulus encoded in a neural population is proportional to the trial-averaged neural spiking rate (Paradiso 1988; Seung & Sompolinsky 1993). If we further assume that each spike has a fixed cost, we find that the local neural cost induced by each item is proportional to *J*,

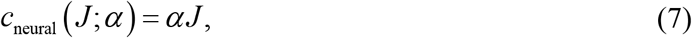

where *α* is the amount of neural cost incurred by a unit increase in resource. Combining Eqs. (5)-(7) yields

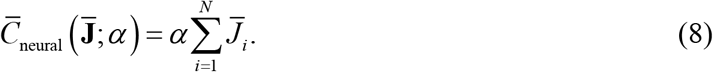

Hence, the global expected neural cost is proportional to the total amount of invested resource and independent of the amount of variability in *J*.

### Formalization of expected behavioral cost for local tasks

Before we specify the expected behavioral cost function, we introduce a distinction between two classes of tasks. First, we define a task as “local” if the observer’s response depends on only one of the encoded items. Examples of local tasks are delayed estimation (Blake et al. 1997; Prinzmetal et al. 1998; Wilken & Ma 2004), single-probe change detection (Todd & Marois 2004; Luck & Vogel 1997), and single-probe change discrimination (Klyszejko et al. 2014). By contrast, when the task response depends on all memorized items, we define the task as “global”. Examples of global tasks are whole-display change detection (Luck & Vogel 1997; Keshvari et al. 2013), change localization (Van Den Berg et al. 2012), and delayed visual search (Mazyar et al. 2012). The theory that we developed up to this point – Eqs. (1) to (8) – applies to both global and local tasks. However, from here on, we developed our theory in the context of local tasks only; we will come back to global tasks at the end of Results.

Since in local tasks only one item gets probed, the total expected behavioral cost is a weighted average of expected costs associated with individual items,

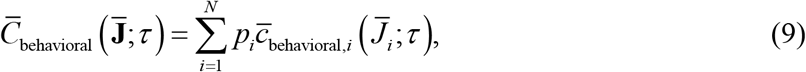

where *p_i_* is the experimentally determined probing probability of the *i*th item and 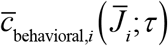 is the local expected behavioral cost associated with reporting the *i*th item. The only remaining step is to specify 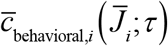. This function is task specific and we will specify it after we have described the task to which we apply the model.

### Formalization of expected behavioral cost for local tasks

Combining Eqs. (3), (8), and (9) yields the following expected total cost function for local tasks:

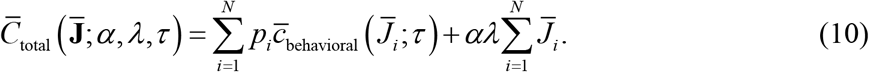

Since parameters *α* and *λ* have interchangeable effects on the model predictions, we will fix *α*=1 and only treat *λ* as a free parameter.

We recognize that the right-hand side of Eq. (10) is a sum of independent “local” expected total costs. Therefore, each element of 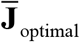, Eq. (4), can be computed independently of the other elements, by minimizing the local expected total cost for each item,

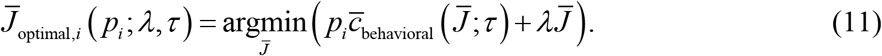

This completes the specification of the general form of our resource-rational model for local tasks. Its free parameters are *λ* and *τ*.

### Set size effects result from cost minimization and are mediated by probing probability

To obtain an understanding of the model predictions, we mathematically analyze how 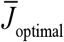 depends on probing probability and set size. We perform this analysis under two very general assumptions about the expected behavioral cost function: first, it monotonically decreases with 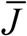 (i.e., increasing resource reduces the expected behavioral cost) and, second, it satisfies a law of diminishing returns (i.e., the reductions per unit increase of resource decrease with the total amount of already invested resource). As proven in the Supplementary Information, under these assumptions the domain of *p_i_* consists of three potential regimes, corresponding to different encoding strategies (Fig. 1A). First, there might exist a regime 0≤*p_i_*<*p*_0_ in which it is optimal to not encode an item, 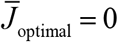. In this regime, the probing probability of an item is so low that investing any amount of resource can never reduce the expected behavioral cost by more than it increases the expected neural cost. Second, there might exist a regime *p*_0_≤*p_i_*<*p*_∞_ in which it is optimal to encode an item with a finite amount of resource, 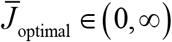. In this regime, 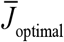 increases as a function of *p_i_*. Finally, there may be a regime *p*_∞_<*p_i_*>1 in which the optimal strategy is to encode the item with an infinite amount of resource, 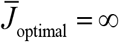. This last regime will only exist in extreme cases, such as when there is no neural cost associated with encoding. The threshold *p*_0_ depends on the importance of the neural cost, *λ*, and on the derivative of the expected behavioral cost evaluated at 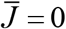; specifically, 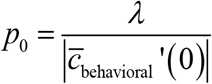. The threshold *p*_∞_ depends on *λ* and on the derivative of the expected behavioral cost evaluated at 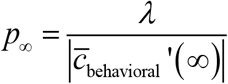; specifically, 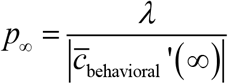.

**Figure 1.**
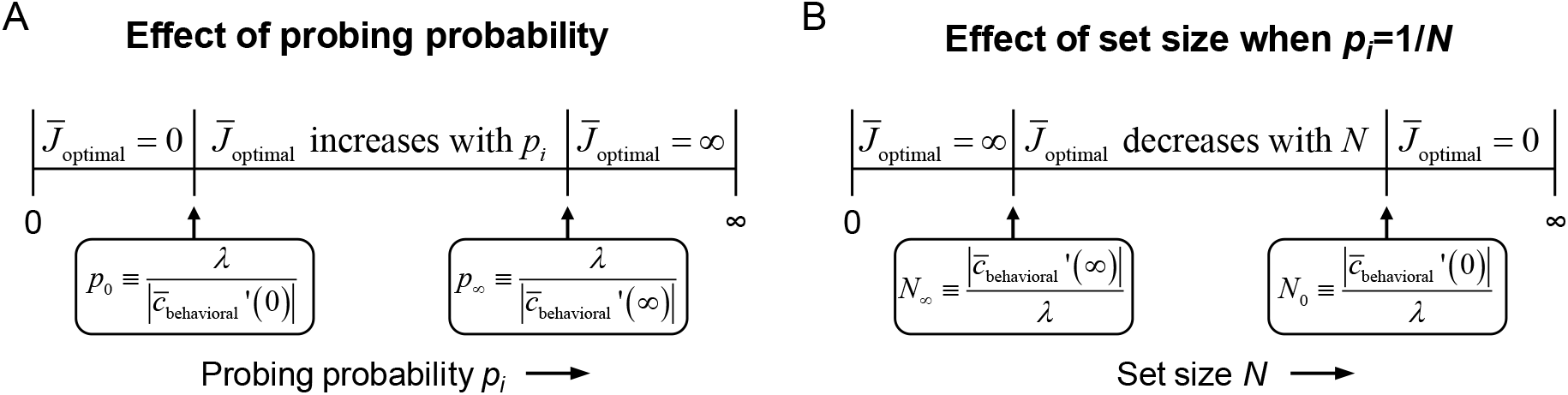
Effects of probing probability and set size on 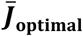 in the resource-rational model for local tasks. (A) The model has three different optimal solutions depending on probing probability *p_i_*: invest no resource when *p_i_* is smaller than some threshold value *p*_0_, invest infinite resource when *p_i_* is larger than *p*_∞_, and invest a finite amount of resource when *p*_0_ < *p_i_* < *p*_∞_. (B) The domain of *N* partitions in a similar manner for tasks in which *p_i_*=1/N.

We next turn to set size effects. An interesting property of the model is that 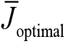 only depends on the probing probability, *p_i_*, and on the model parameters – it does *not* explicitly depend on set size, *N*. Therefore, the only way that it can predict set size effects is through a coupling between *N* and *p_i_*. Such a coupling exists in most studies that use a local task. For example, in delayed-estimation tasks, each item is usually equally likely to be probed such that *p_i_*=1/*N*. For those experiments, the above partitioning of the domain of *p_i_* translates to a similar partitioning of the domain of *N* (Fig. 1B). Then, a set size *N*_∞_≥0 may exist below which it is optimal to encode items with infinite resource, a region *N*_∞_≤*N*<*N*_0_ in which it is optimal to encode items with a finite amount of resource, and a region *N*>*N*_0_ in which it is optimal to not encode items at all.

## RESULTS

### Model predictions for delayed estimation tasks

To test the predictions of the model against empirical data, we apply it to the delayed estimation task (Wilken & Ma 2004; Blake et al. 1997; Prinzmetal et al. 1998), which is currently one of the most widely used paradigms in VWM research. In this task, the observer briefly holds a set of items in memory and then reports their estimate of a randomly probed target item (Fig. 2A). Set size effects manifest as a widening of the estimation error distribution as the number of items is increased (Fig. 2B), which suggests a decrease in the amount of resource per item (Fig. 2C).

**Figure 2.**
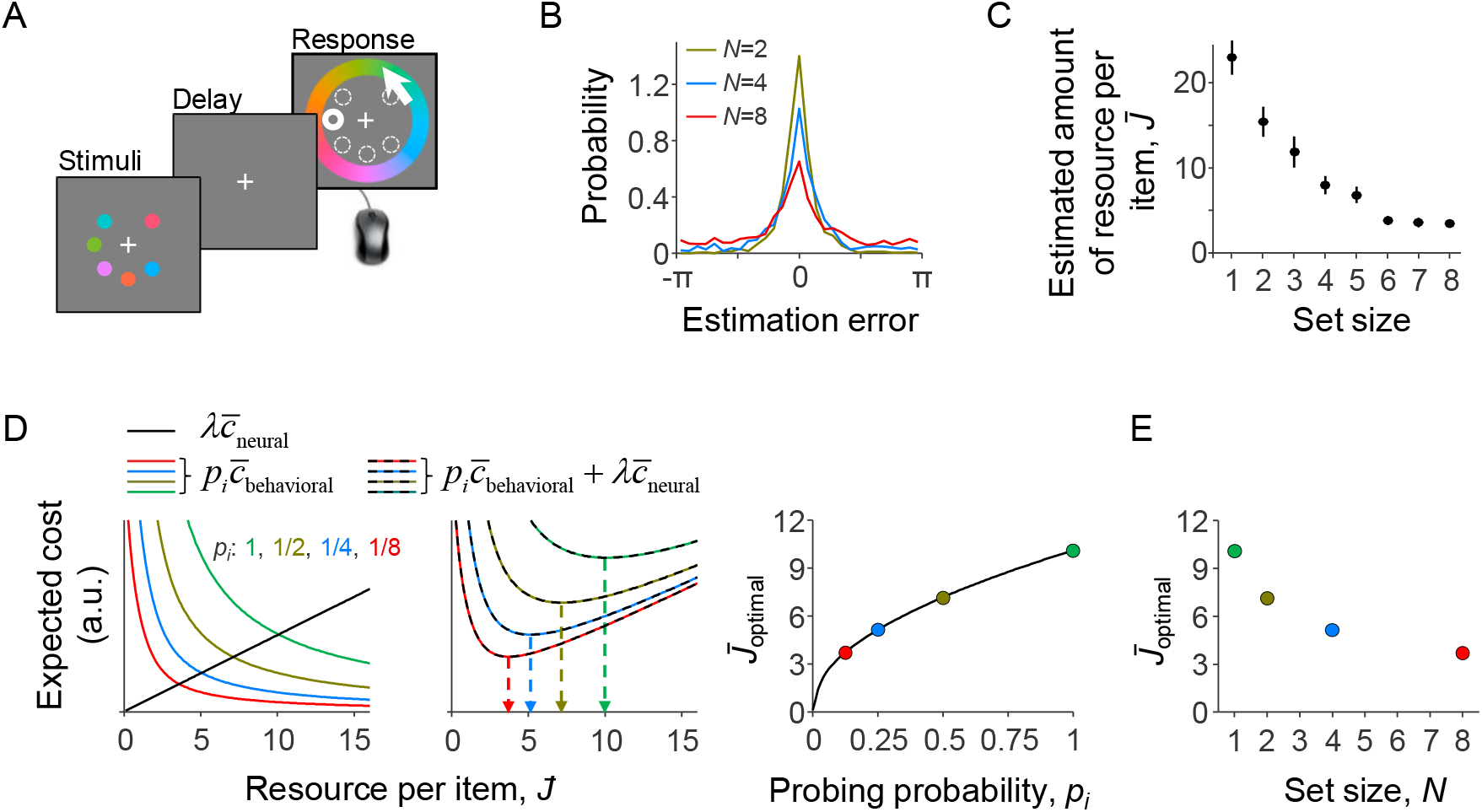
A resource-rational model for delayed-estimation tasks. (A) Example of a trial in a delayed-estimation experiment. The subject is briefly presented with a set of stimuli and, after a short delay, reports the value of the item at a randomly chosen location (here indicated with thick circle). (B) The distribution of estimation errors in delayed-estimation experiments typically widens with set size (data from Experiment E5 in Table 1). (C) This set size effect can be explained as a decrease in the amount of resource per encoded item. The estimated amount of resource per item was computed using the same non-parameter model as in 3C. (D) Expected cost under four different probing probabilities (model parameters: *λ*=0.001, *β*=2, τ↓0). *Left*: The local expected behavioral cost multiplied by *p_i_* (colored curves) decreases with the amount of invested resource, while the expected neural cost increases (black line). *Center*: The sum of these two costs has a unique minimum, whose location (arrows) depends on probing probability *p_i_*. *Right*: The optimal amount of resource per item increases with the probability that the item will be probed. (E) The optimal amount of resource per item from panel *C* replotted as a function of set size, *N*, for a task in which all items are equally likely to be probed, i.e., *p_i_*=1/*N*. The predicted set size effect is qualitatively similar to set size effects observed in empirical data (cf. panel C).

To apply our model to this task, we express the expected local behavioral cost as an expected value of a local behavioral cost with respect to the error distribution,

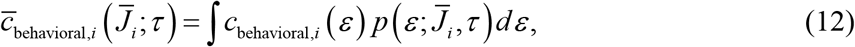

where the behavioral cost function *c*_behavioral,*i*_(*ε*) maps an encoding error *ε* to a cost and 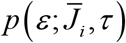 is the predicted distribution of *ε* for an item encoded with resource 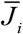. We first specify 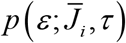 and then turn to *c*_behavioral,*i*_(*ε*). Since the task-relevant feature in delayed-estimation experiments is usually a circular variable (color or orientation), we make the common assumption that *ε* follows a Von Mises distribution. We denote this distribution by VM(*ε*;*J*), where *J* is one-to-one related to the distribution’s concentration parameter *κ* (see Supplementary Information). The distribution of *ε* for a stimulus encoded with resource 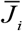. is found by integrating over *J*,

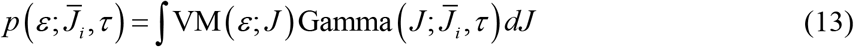

Finally, we specify the behavioral cost function *c*_behavioral,*i*_ (*ε*) in Eq. (12), which maps an estimation error *ε* to a behavioral cost. As in most psychophysical experiments, human subjects tend to perform well on delayed-estimation tasks even when the reward is independent of their performance. This suggests that the behavioral cost function is strongly determined by internally incentives. A recent paper (Sims 2015) has attempted to measure this mapping and proposed a two-parameter function. We will test that proposal later, but for the moment we assume a simpler, one-parameter power-law function, *c*_behavioral,*i*_ (*ε*; *β*) = |*ε*|^*β*^, where power *β* is a free parameter.

To get an intuition for the predictions of this model, we plot the expected behavioral cost, the expected neural cost, and their sum as a function of 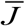 for a specific set of parameters and for four different values of probing probability *p_i_* (Fig. 2D). The expected total cost has a unique minimum in all four cases and the value of 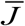 corresponding to this minimum increases with *p_i_*. Hence, in this example, the optimal amount of resource assigned to an item is an increasing function of its probing probability.

The probing probabilities in Fig. 2D correspond to the probing probabilities of items at set sizes 1, 2, 4, and 8 in a task where each item is equally likely to be probed, *p_i_*=1/*N*. When we replot the values of 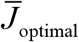 from Fig. 2D as a function of set size, we observe a set size effect that is qualitatively similar to effects observed in empirical data (Fig. 2E; cf. Fig. 2C). Hence, the model predicts set size effects in delayed estimation tasks, even though these effects are fully mediated by individual-item probing probability. This notion is reminiscent of empirical observations. Palmer (1993) reported that “relevant set size” (where irrelevance means *p_i_*=0) acts virtually identically to actual set size. Emrich et al. (2017) independently varied probing probability and set size in their experiment, and found that the former was a better predictor of performance than the latter. Based on this, they hypothesized that set size effects are mediated by probing probability. These findings are – at least qualitatively – consistent with the predictions of our model.

### Model fits to data from delayed-estimation experiments with equal probing probabilities

To examine how well the model accounts for set size effects in empirical data, we fit it to data from six experiments that are part of a previously published benchmark set (E1-E6 in Table 1)^*^. We use a Bayesian optimization method (Acerbi & Ma 2017) to estimate the maximum-likelihood parameter values, separately for each individual data set (see Supplementary Table S1 for a summary of the estimates). The results show that the model accounts well for the subject-level error distributions (Fig. 3A) and the two statistics that summarize these distributions (Fig. 3B).

**Table 1.** Overview of experimental datasets. Experiments E5 and E6 differed in the way that subjects provided their responses (E5: color wheel; E6: scroll).

| ID | Reference | Feature | Set size(s) | Probing probability | Number of subjects |
| --- | --- | --- | --- | --- | --- |
| E1 | (Wilken & Ma 2004) | Color | 1, 2, 4, 8 | Equal | 15 |
| E2 | (Zhang & Luck 2008) | Color | 1, 2, 3, 6 | Equal | 8 |
| E3 | (Bays et al. 2009) | Color | 1, 2, 4, 6 | Equal | 12 |
| E4 | (Van Den Berg et al. 2012) | Orientation | 1-8 | Equal | 6 |
| E5 | (Van Den Berg et al. 2012) | Color | 1-8 | Equal | 13 |
| E6 | (Van Den Berg et al. 2012) | Color | 1-8 | Equal | 13 |
| E7 | (Bays 2014) | Orientation | 2,4,8 | Unequal | 7 |
| E8 | (Emrich et al. 2017) | Color | 4 | Unequal | 20 |
| E9 | (Emrich et al. 2017) | Color | 6 | Unequal | 20 |

**Figure 3.**
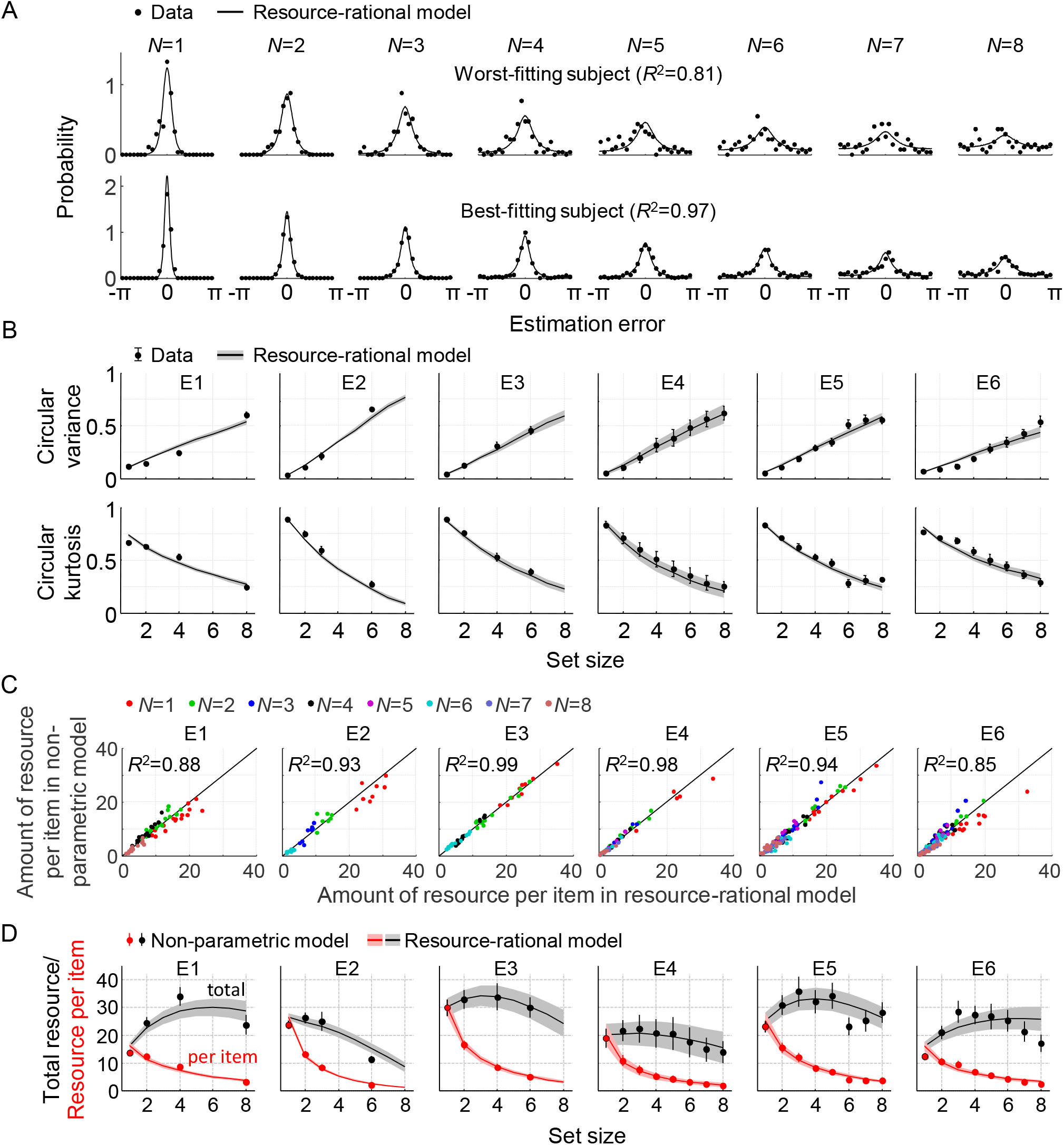
Model fits to data from six delayed-estimation experiments with equal probing probabilities. (A) Maximum-likelihood fits to raw data of the worst-fitting and best-fitting subjects. Goodness of fit was measured as *R*^2^, computed for each subject by concatenating histograms across set sizes. (B) Subject-averaged circular variance and kurtosis of the estimation error, as a function of set size and split by experiment. The maximum-likelihood fits of the model account well for the trends in these statistics. (C) Estimated amounts of resource per item in the resource-rational model scattered against the estimates in the non-parametric model. Each dot represents estimates from a single subject. (D) Estimated amount of resource per item (red) and total resource (black) plotted against set size. Here and in subsequent figures, error bars and shaded areas represent 1 s.e.m. of the mean across subjects.

We next compare the goodness of fit of the resource-rational model to that of a descriptive variant in which the amount of resource per item, 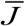, is assumed to be a power-law function of set size (all other aspects of the model are kept the same). This variant is identical to the VP-A model in our earlier work (van den Berg et al. 2014). Model comparison based on the Akaike Information Criterion (AIC) (Akaike 1974) indicates that the data provide similar support for both models, with a small advantage for the resource-rational model (ΔAIC=5.27±0.70; throughout the paper, X±Y indicates mean±s.e.m. across subjects). Hence, the resource-rational model provides a principled explanation of set size effects without sacrificing quality of fit compared to one of the best available descriptive models of VWM^†^. We find that the resource-rational model also fits better than a model in which the total amount of resource is fixed and divided equally across items (ΔAIC=13.9±1.4).

So far, we have assumed that there is random variability in the actual amount of resource assigned to an item. Next, we test an equal-precision variant of the resource-rational model, by fixing parameter *τ* to a very small value (10^−3^). Consistent with the results obtained with the variable-precision model, we find that the rational model has a substantial AIC advantage over a fixed-resource model (ΔAIC=43.0±6.8) and is at equal footing with the power-law model (ΔAIC=2.0±1.7 in favor of the power-law model). However, all three equal-precision models (fixed resource, power law, rational) are outperformed by their variable-precision equivalents by over 100 AIC points. Therefore, we will only consider variable-precision models in the remainder of the paper.

To get an indication of the absolute goodness of fit of the resource-rational model, we next examine how much room for improvement there is in the fits. We do this by fitting a non-parametric model variant in which resource 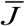 is a free parameter at each set size, while keeping all other aspects of the model the same. We find a marginal AIC difference, which indicates that the fits of the rational model cannot be improved much further without overfitting the data (ΔAIC=3.49±0.93, in favor of the non-parametric model). An examination of the fitted parameter values corroborates this finding: the estimated resource values in the non-parametric model closely match the optimal values in the rational model (Fig. 3C).

So far, we have assumed that behavioral cost is a power-law function of the absolute estimation error, *c*_behavioral_(*ε*)=|*ε*|^*β*^. To evaluate the necessity of a free parameter in this function, we also test three parameter-free choices: |*ε*|, *ε*^2^, and –cos(*ε*). Model comparison favors the original model with AIC differences of 14.0±2.8, 24.4±4.1, and 19.5±3.5, respectively. While there may be other parameter-free functions that give better fits, we expect that a free parameter is unavoidable here, as the error-to-cost mapping may differ across experiments (due to differences in external incentives) and also across subjects within an experiment (due to differences in intrinsic motivation). Finally, we also test a two-parameter function that was proposed recently (Eq. (5) in (Sims 2015)). The main difference with our original choice is that this alternative function allows for saturation effects in the error-to-cost mapping. However, this extra flexibility does not increase the goodness of fit sufficiently to justify the additional parameter, as the original model outperforms this variant with an AIC difference of 5.3±1.8.

Finally, we use five-fold cross validation to verify the AIC-based results reported in this section. We find that they are all consistent (Table S2 in Supplementary Information).

### Non-monotonic relation between total resource and set size

One quantitative feature that sets the resource-rational theory apart from previous theories is its predicted relation between set size and the total amount of invested resource, 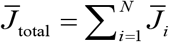. This quantity is – by definition – constant in fixed-resource models, and in power-law models it varies monotonically with set size. By contrast, we find that in the fits to several of the experiments, 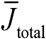 varies *non-monotonically* with set size (Fig. 3D, gray curves). To examine whether there is evidence for non-monotonic trends in the subject data, we next compute an “empirical” estimate 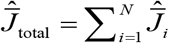, where 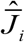 are the best-fitting resource estimates in the non-parametric model. We find that these estimates show evidence of similar non-monotonic relations in some of the experiments (Fig. 3D, black circles). To quantify this evidence, we perform Bayesian paired t-tests in which we compare the estimates of 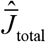 at set size 3 with the estimates at set sizes 1 and 6 in the experiments that included these three set sizes (E2 and E4-E6). These tests reveal strong evidence that the total amount of resource is higher at set size 3 than at set sizes 1 (BF_+0_=1.05·10^7^) and 6 (BF_+0_=4.02·10^2^). We next compute for each subject the set size at which 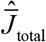 is largest, which we denote by *N*_peak_, and find a subject-averaged value of 3.52±0.18. Altogether, these findings suggest that the total amount of resource that subjects spend on item encoding varies non-monotonically with set size, which is consistent with predictions from the resource-rational model, but not with any of the previously proposed models. To the best of our knowledge, evidence for a possible non-monotonicity in the relation between set size and total encoding resource has not been reported before.

### Predicted effects of probing probability

As we noted before, the model predictions do not explicitly depend on set size, *N*. Yet, we found that the model accounts well for set size effects in the experiments that we considered so far (E1-E6). The resolution of this paradox is that in all those experiments, *N* was directly coupled with probing probability *p_i_*, through *p_i_*=1/*N*. This coupling makes it impossible to determine whether changes in subjects’ encoding precision are due to changes in *N* or due to changes in *p_i_*. Therefore, we will next consider experiments in which individual probing probabilities and set size were varied independently of each other (E7-E9 in Table 1). According to our model, the effects of *N* that we found in E1-E6 were really effects of *p_i_*. Therefore, we should be able to make predictions about effects of *p_i_* in E7-E9 by recasting the effects of *N* in E1-E6 as effects of *p_i_*=1/*N*. Given that the amount of resource per item in E1-E6 decreases with *N*, a first prediction is that it should increase as a function of *p_i_* in E7-E9. A second and particularly interesting prediction is that the estimated total amount of invested resource should vary non-monotonically with *p_i_* and peak at a value *p*_peak_ that is close to 1/*N*_peak_ found in E1-E6 (see previous section). Based on the values of *N*_peak_ in experiments E1-E6, we find a prediction *p*_peak_=0.358±0.026.

### Model fits to data from delayed-estimation experiments with unequal probing probabilities

To test the predictions presented in the previous section and, more generally, to evaluate how well our model accounts for effects of *p_i_* on encoding precision, we fit it to data from three experiments in which probing probability was varied independently of set size (E7-E9 in Table 1).

In the first of these experiments (E7), seven subjects performed a delayed-estimation task at set sizes 2, 4, and 8. On each trial, one of the items – indicated with a cue – was three times more likely to be probed than any of the other items. Hence, the probing probabilities for the cued and uncued items were 3/4 and 1/4 at *N*=2, respectively, 1/2 and 1/6 at *N*=4, and 3/10 and 1/10 at *N*=8. The subject data show a clear effect of *p_i_*: the higher the probing probability of an item, the more precise the subject responses (Fig. 4A, top row, black circles). We find that the resource-rational model, Eq. (11), accounts well for this effect (Fig. 4A, top row, curves) and does so by increasing the amount of resource as a function of probing probability *p_i_* (Fig. 4B, left panel, red curves).

In the other two experiments (E8 and E9), the number of cued items and cue validity were varied between conditions, while set size was kept constant at 4 or 6. For example, in one of the conditions of E8, three of the four items were cued with 100% validity, such that *p_i_* was 1/3 for each cued item and 0 for the uncued item; in another condition of the same experiment, two of the four items were cued with 66.7% validity, meaning that *p_i_* was 1/3 for each cued item and 1/6 for each uncued item. The unique values of *p_i_* across all conditions were {0, 1/6, 2/9, 1/4, 1/3, 1/2, 1} in E8 and {0, 1/12, 1/10, 2/15, 1/6, 1/3, 1/2, and 1} in E9. As in E7, responses get more precise with increasing *p_i_* and the model accounts well for this (Fig. 4A), again by increasing the amount of resource assigned to an item with *p_i_* (Fig. 4B).

We next examine how our model compares to the models proposed in the papers that originally published these three datasets. In contrast to our model, both Bays (2014) and Emrich et al. (2017) proposed that the total amount of invested resource is fixed. However, while Bays proposed that the distribution of this resource is in accordance with minimization of a behavioral cost function (as in our model), Emrich et al. postulated that the resource is distributed in proportion to each item’s probing probability. Hence, while our model optimizes both the amount of invested resource and its distribution, Bays’ model only optimizes the distribution, and Emrich et al.’s model does not explicitly optimize anything. To examine how the three proposals compare in terms of how well they account for the data, we fit two variants of our model that encapsulate the main assumptions of these two earlier proposals. In the first variant, we compute 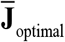 as 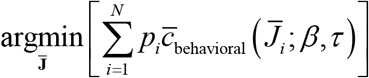 under the constraint 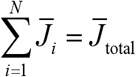, which is consistent with Bays’s proposal. Hence, in this variant, the neural cost function is removed and parameter *λ* is replaced by a parameter 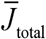 – otherwise, all aspects of the model are the same as in our main model. In the variant that we use to test Emrich et al.’s proposal, we compute 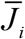 for each item as 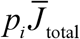, where *p_i_* is the probing probability and 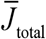 is again a free parameter that represents the total amount of resource. Comparing the models using the data from all 47 subjects of E7-E9, we find a substantial advantage of our model over the proposal by Emrich et al., with an AIC difference of 18.0±3.9. However, our model cannot reliably be distinguished from the proposal by Bays: both models are preferred in about half of the subjects (our model: 27; Bays: 20) and the subject-averaged AIC difference is negligible (1.8±2.5 in favor of our model). Hence, the model comparison suggests quite convincingly that subjects distribute their resource near-optimally across items with unequal probing probabilities, but it is inconclusive regarding the question whether the total amount of invested resource is fixed or optimized.

As an alternative way to address the question of whether the total amount of resource is fixed, we again fit a non-parametric model to obtain “empirical” estimates of the total amount of invested resource. To this end, we define 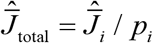, where 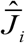 are the best-fitting values in a non-parametric model, such that 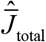 represents the estimated total amount of resource that a subject would invest to encode a display filled with items that all have probing probability *p_i_*. We find that these estimates show signs of a non-monotonicity as a function of *p_i_* (Fig. 4B, black points), which are captured reasonably well by the resource-rational model (Fig. 4B, black curves). Averaged across all subjects in E7-E9, the value of *p_i_* at which 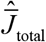 is largest is 0.384±0.037, which is close to the predicted value of 0.358±0.026 (see previous section). Indeed, a Bayesian independent-samples t-test supports the null hypothesis that there is no difference (BF01=4.27). Hence, while the model comparison results in the previous paragraph were inconclusive regarding the question whether the total amount of invested resource is fixed or optimized, the present analysis provides evidence against fixed-resource models and confirms a prediction made by our own model.

In summary, the results in this section show that effects of probing probability in E7-E9 are well accounted for by the same model as we used to explain effects of set size in E1-E6. Regardless of whether total resource is fixed or optimized, this finding provides further support for the suggestion that set size effects are mediated by probing probability (Emrich et al. 2017) or, more generally, by item relevance (Palmer et al. 1993).

### Is it ever optimal to not encode an item?

There is an ongoing debate about the question whether a task-relevant item is sometimes completely left out of working memory (Adam et al. 2017; Luck & Vogel 2013; Ma et al. 2014; Rouder et al. 2008). Specifically, slot models predict that this happens when set size exceeds the number of slots (Zhang & Luck 2008). In resource models, the possibility of complete forgetting has so far been an added ingredient separate from the core of the model (van den Berg et al. 2014). Our normative theory allows for a reinterpretation of the question: are there situations in which it is optimal to assign zero resource to the encoding of an item? We already established that this could happen in delayed-estimation tasks: whenever the probing probability is lower than a threshold value 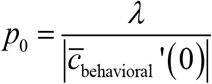, the optimal amount of resource to invest on encoding the item is zero (see Theory). But what values does *p*_0_ take in practice? Considering the expected behavioral cost function of a fixed-precision model (a variable-precision model with *τ* ↓ 0), we can prove that *p*_0_=0, i.e. it is never optimal to not invest any resource (Supplementary Information). For the expected behavioral cost function of the general variable-precision model, however, simulations indicate that *p*_0_ can be greater than 0 (we were not able to derive this result analytically). We next examine whether this ever happens under parameter values that are representative for human subjects. Using the maximum-likelihood parameters obtained from the data in E7-E9, we estimate that *p*_0_ (expressed as a percentage) equals 8.86 ± 0.54%. Moreover, we find that for 8 of the 47 subjects, *p*_0_ is larger than the lowest probing probability in the experiment, which suggests that these subjects sometimes entirely ignored one or more of the items. For these subjects, the error distributions on items with *p_i_*<*p*_0_ look uniform (see Fig. 4C for an example) and Kolmogorov-Smirnov tests for uniformity did not reject the null hypothesis in any of these cases (*p*>0.05 in all tests).

These results suggest that their might be a principled reason for why people sometimes leave task-relevant items out of visual working memory in delayed-estimation experiments. However, our model cannot explain all previously reported evidence for this. In particular, when probing probabilities are equal for all items, the model makes an “all or none” prediction: all items are encoded when *p_i_*>*p*_0_ and none otherwise. Hence, it cannot explain why subjects in tasks with equal probing probabilities sometimes seem to encode a subset of task-relevant items. For example, a recent study reported that in a whole-report delayed-estimation experiment (*p_i_*=1 for all items), subjects encoded about half of the 6 presented items on each trial (Adam et al. 2017). Unless additional assumptions are made, our model cannot account for this finding.

### Predictions for a global task: whole-display change detection

The results so far show that the resource-rational model accounts well for data in a variety of delayed-estimation experiments. To examine how its predictions generalize to other tasks, we next consider a change detection task, which is another widely used paradigm in research on VWM. In this task, the observer is sequentially presented with two sets of items and reports if any one of them changed (Fig. 5A). In the variant that we consider here, a change is present on exactly half of the trials and is equally likely to occur in any of the items. We construct a model for this task by combining Eqs. (3), (4), and (8) with an expected behavioral cost function based on the Bayesian decision rule for this task (see Supplementary Information), which yields

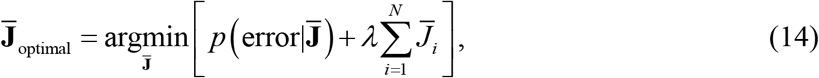

where 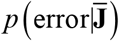 is the expected behavioral cost function, which in this case specifies the probability of an error response when a set of items is encoded with resource 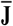.

In contrast to local tasks, the total expected cost in global tasks cannot be written as a sum of local expected costs, because the expected behavioral cost – such as 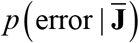 in Eq. (14) – can only be computed globally, not per item. Consequently, the elements of 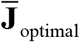 in global tasks cannot be computed separately for each item. This makes resource optimization computationally much more demanding, because it requires solving an *N*-dimensional minimization problem instead of *N* one-dimensional problems.

We next perform a simulation at *N*=2 (which is still tractable) to get an intuition of the predictions that follow from Eq. (14). For practical convenience, we assume in this simulation that there is no variability in precision, *τ*↓0, such that *λ* is the only model parameter. The results (Fig. 5B) show that the cost-minimizing strategy is to encode neither of the items when the amount of reward per correct trial is very low (left panel) and encode them both when reward is high (right panel). However, interestingly, there is also an intermediate regime in which the optimal strategy is to encode one of the two items, but not the other one (Fig. 5B, central panel). Hence, just as in the delayed-estimation task, there are conditions in which it is optimal to encode only a subset of items. An important difference, however, is that in the delayed-estimation task this only happens when items have unequal probing probabilities, while in this change detection task it even happens when all items are equally likely to change.

Simulations at larger set sizes quickly become computationally intractable, because of the reason mentioned above. However, the results at *N*=2 suggest that if two items are encoded, the optimal solution is to encode them with the same amount of resource (Fig. 5C). Therefore, we conjecture that all non-zero values in 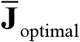 are identical, which would mean that the entire vector can be summarized by two values: the number of encoded items, which we denote by *K*_optimal_, and the amount of resource assigned to each encoded item, which we denote by 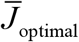. Using this conjecture (which we have not yet been able to prove), we are able to efficiently compute predictions at an arbitrary set size. Simulation results show that the model then predicts that both *K*_optimal_ and 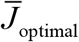 depend on set size (Fig 4D, left) and produces response data that are qualitatively similar to human data (Fig. 5D, right).

**Figure 4.**
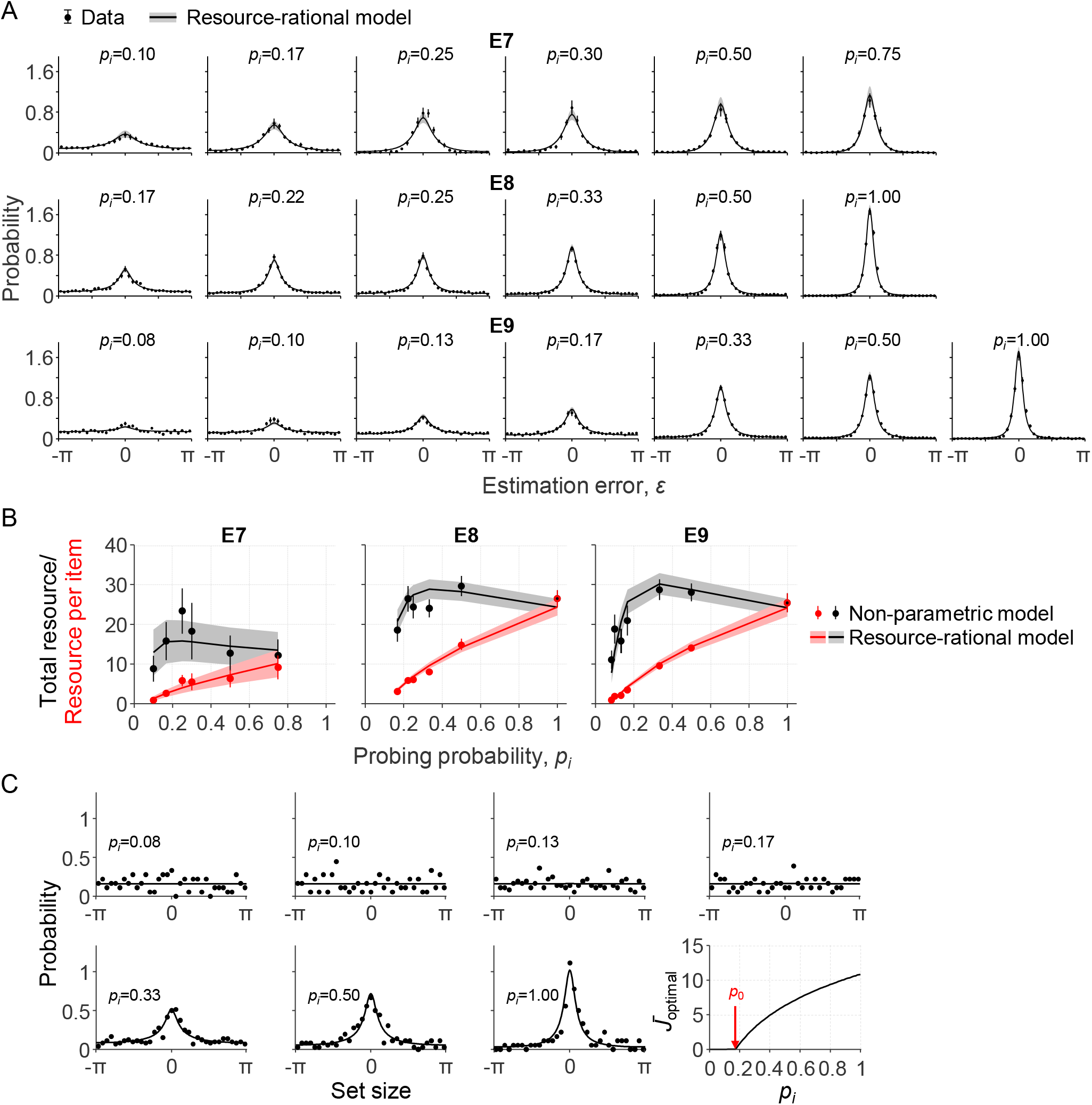
Model fits to data from three delayed-estimation experiments with unequal probing probabilities. (A) Fits of the resource-rational model (curves) to the data (black circles) of experiments E7-E9. (B) Estimated amount of resource per item as a function of probing probability (red) and the corresponding estimated total amount of resource that the subject would spend on encoding a display filled with items with the equal probing probabilities (black). (C) Error histograms and a plot of *J*_optimal_ as a function of *p_i_* for one of the subjects with an estimated value of *p*_0_ larger than the smallest probing probability (subject S4 in E9; *p*_0_=0.18). The error histograms for items with the four lowest probing probabilities appear to be uniform for this subject, which is indicative of guessing (*p*>0.23 in Kolgomorov-Smirnov tests for uniformity on these four distributions).

**Figure 5.**
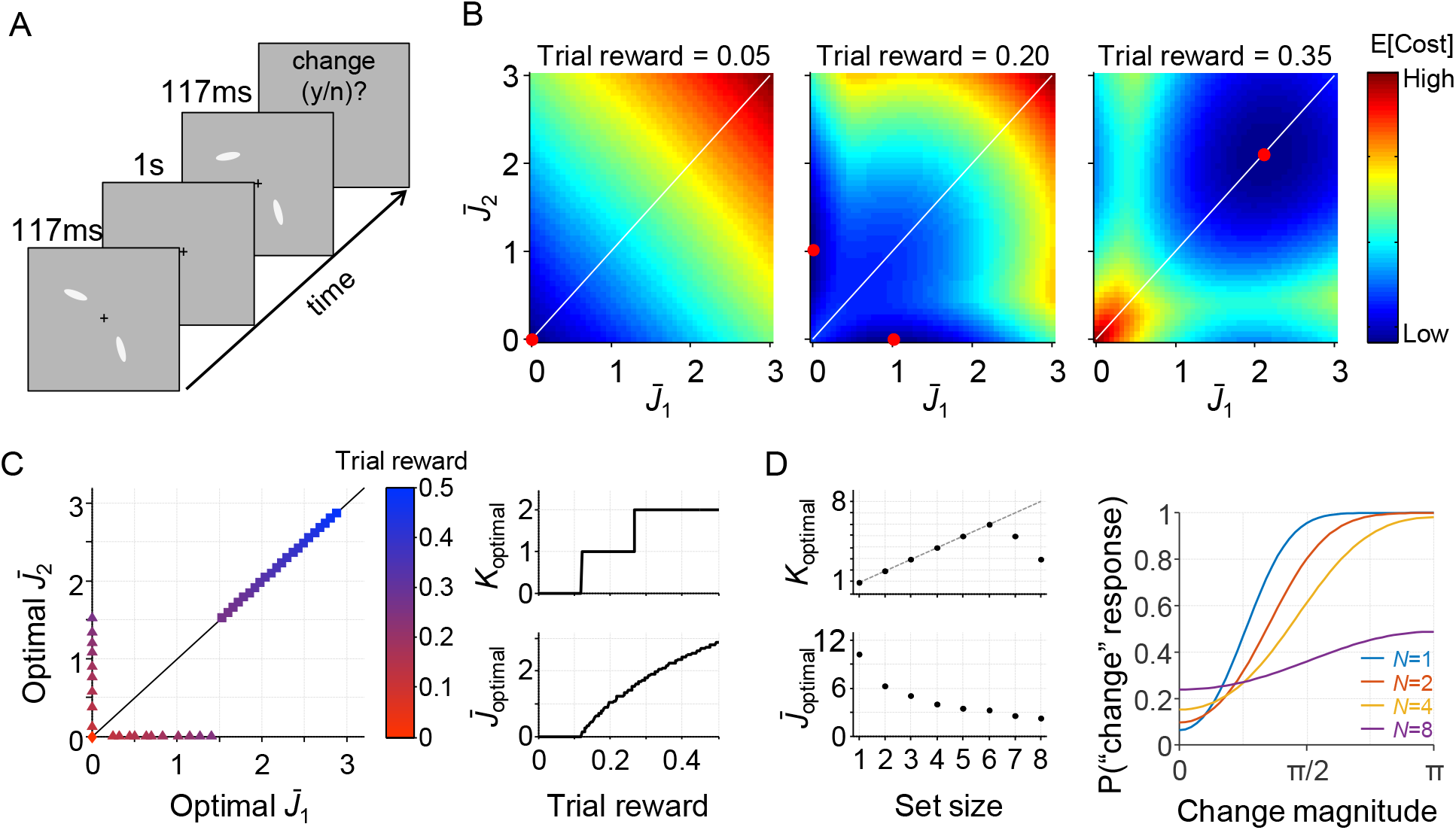
A resource-rational model for change-detection tasks. (A) Example of a trial in a change detection task with a set size of 2. The subject is sequentially presented with two sets of stimuli and reports whether there was a change at any of the item locations. (B) Simulated expected total cost in the resource-rational cost function applied to a task with a set size of 2 an a reward of 0.05 (left), 0.20 (center), or 0.35 (right) units per correct trial. The red dot indicates the location of minimum cost, i.e., the resource-optimal combination of 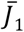 and 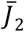 (note that the expected cost function in the central panel has a minimum at two distinct locations). When reward is low (left), the optimal strategy is to encode neither of the two stimuli. When reward is high (right), the optimal strategy is to encode both stimuli with equal amounts of resource. For intermediate reward (center), the optimal strategy is to encode one of the two items, but not the other one. (C) Model predictions as a function of trial rewards at *N*=2. Left: The amount of resource assigned to the two items for a range of reward values. Right: the corresponding optimal number of encoded items (top) and optimal amount of resource per encoded item (bottom) as a function of reward. (D) Model predictions as a function of set size (trial reward = 1.5). The model predicts set size effects in both the number of encoded items (left, top) and the amount of resource with which these items are encoded (left, bottom). Moreover, the model produces response data (right) that are qualitatively similar to human data (e.g., Keshvari et al. 2013). The parameter values used in all simulations were *λ*=0.01 and τ↓0.

## DISCUSSION

### Summary

Descriptive models of visual working memory (VWM) have evolved to a point where there is little room for improvement in how well they account for experimental data. Nevertheless, the basic finding that VWM precision depends on set size still lacks a principled explanation. Here, we examined a normative proposal in which expected task performance is traded off against the cost of spending neural resource on encoding. We used this principle to construct a resource-rational model for “local” VWM tasks and found that set size effects in this model are fully mediated by the probing probabilities of the individual items; this is consistent with suggestions from earlier empirical work (Emrich et al. 2017; Palmer et al. 1993). From the perspective of our model, the interpretation is that as more items are added to a task, the relevance of each individual item decreases, which makes it less cost-efficient to spend resource on its encoding. We also found that in this model it is sometimes optimal to encode only a subset of task-relevant items, which implies that resource rationality could serve as a principled bridge between resource and slot-based models of visual working memory. We tested the model on data from nine previous delayed-estimation experiments and found that it accounts well for effects of both set size and probing probability, despite having relatively few parameters. Moreover, it accounts for a non-monotonicity that appears to exist between set size and the total amount of resource that subjects invest in item encoding. The broader implication of our findings is that VWM limitations – and cognitive limitations in general – may be driven by a mechanism that minimizes a cost, instead of by a fixed constraint on available encoding resource.

### Limitations

Our theory makes a number of assumptions that need further investigation. First, we have assumed that the expected behavioral cost decreases indefinitely with the amount of invested resource, such that in the limit of infinite resource there is no encoding error and no behavioral cost. However, encoding precision in VWM is fundamentally limited by the precision of the sensory input, which is itself limited by irreducible sources of neural noise – such as Johnson noise and Poisson shot noise (Faisal et al. 2008; Smith 2015) – and suboptimalities in early sensory processing (Beck et al. 2012). One way to incorporate this limitation is by assuming that there is a resource value 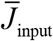 beyond which the expected behavioral cost no longer decreases as a function of 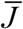. In this variant, 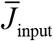 represents the quality of the input and 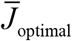 will never exceed this value, because any additional resource would increase the expected neural cost without decreasing the expected behavioral cost.

Second, our theory assumes that there is no upper limit on the total amount of resource available for encoding: cost is the only factor that matters. However, since the brain is a finite entity, the total amount of resource must obviously have an upper limit. This constraint can be incorporated by optimizing **J**_optimal_ under the constraint 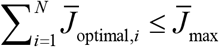, where 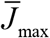 represents the maximum amount of resource that can be invested. While an upper limit certainly exists, it may be much higher than the average amount of resource needed to encode information with the same fidelity as the sensory input. If that is the case, then 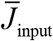 would be the constraining factor and 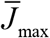 would have no effect.

Third, our theory assumes that there is no lower limit on the amount of resource available for encoding. However, there is evidence that task-irrelevant stimuli are sometimes automatically encoded (Yi et al. 2004; Shin & Ma 2016), perhaps because in natural environments few stimuli are ever completely irrelevant. This would mean that there is a lower limit to the amount of resource spent on encoding. In contradiction to the predictions of our model, such a lower limit would prevent subjects from sometimes encoding nothing at all. For local tasks, such a lower limit can be incorporated by assuming that probing probability *p_i_* is never zero.

We have fitted our model only to data from delayed-estimation experiments. However, it applies without modification to other local tasks, such as single-probe change detection (Luck & Vogel 1997; Todd & Marois 2004) and single-probe change discrimination (Klyszejko et al. 2014). Further work is needed to examine how well the model accounts for empirical data of such tasks. Moreover, it should be further examine how the theory generalizes to global tasks. One such task could be whole-report change detection; we presented simulation results for this task but the theory remains to be further worked out and fitted to the data.

A final limitation is that our theory assumes that items are uniformly distributed and uncorrelated. Although this is correct for most experimental settings, items in more naturalistic settings are often correlated and can take non-uniform distributions. In such environments, the expected total cost can probably be further minimized by taking into account statistical regularities (Orhan et al. 2014). Moreover, recent work has suggested that even when items are uncorrelated and uniformly distributed, the expected estimation error can sometimes be reduced by using a “chunking”strategy, i.e., encoding similar items as one (Nassar et al. 2018). However, since Nassar et al. assumed a fixed total resource and did not take neural encoding cost into account in their optimization, it remains to be seen whether chunking is also optimal in the kind of model that we proposed. We speculate that this is likely to be the case, because encoding multiple items as one will reduce the expected neural cost (fewer items to encode), while the increase in expected behavioral cost will be negligible if the items are very similar. Hence, it seems worthwhile to examine models that combine resource rationality with chunking.

### Variability in resource assignment

Throughout the paper, we have assumed that there is variability in resource assignment. Part of this variability is possibly due to stochastic factors, but part of it may also be systematic – for example, particular colors and orientations may be encoded with higher precision than others (Bae et al. 2014; Girshick et al. 2011). Whereas the systematic component could have a rational basis (e.g., higher precision for colors and orientations that occur more frequently in natural scenes (Ganguli & Simoncelli 2010; Wei & Stocker 2015)), this is unlikely to be true for the random component. Indeed, when we jointly optimize 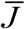 and *τ* in Eq. (11), we find estimates of *τ* that consistently approach 0, meaning that any variability in encoding precision is suboptimal under our proposed cost function. One way to reconcile this apparent suboptimality with the otherwise normative theory is to postulate that maintaining exactly equal resource assignment across cortical regions may itself be a costly process; under such a cost, it could be optimal to allow for some variability in resource assignment. Another possibility is that there are unavoidable imperfections in mental inference (Drugowitsch et al. 2016) that make it impossible to compute 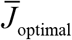 without error, such that the outcome of the computation will vary from trial to trial even when the stimuli are identical.

### Experimental predictions of incentive manipulations

In the present study, we have focused on effects of set size and probing probability on encoding precision. However, our theory also makes predictions about effects of incentive manipulations on encoding precision, because such manipulations affect the expected behavioral cost function.

Incentives can be experimentally manipulated in a variety of ways. One method that was used in at least two previously published delayed-estimation experiments is to make the feedback binary (“correct”, “error”) and vary the value of the maximum error allowed to receive positive feedback (Zhang & Luck 2011; Nassar et al. 2018). In both studies, subjects in a “low precision” condition received positive feedback whenever their estimation error was smaller than a threshold value of π/3. Subjects in the “high precision” condition, however, received positive feedback only when the error was smaller than π/12 (Zhang & Luck 2011) or π/8 (Nassar et al. 2018). In our model, this manipulation can be implemented as a behavioral cost function *c*_behavioral,*i*_(*ε*) that maps values of |*ε*| smaller than the feedback threshold (π/3, π/8, π/12) to 0 and larger values to 1. Neither of the two studies found evidence for a difference in encoding precision between the low- and high-precision conditions. At first, this may seem to be at odds with the predictions of our model, as one may expect that it should assign more resource to items in the high-precision condition. However, simulation results show that the model predictions are not straightforward and that it can account for the absence of an effect (Fig. S4 in Supplementary Information). In particular, the simulation results suggest that the experimental manipulations in the studies by Zhang & Luck and Nassar et al. may not have been strong enough to measure an effect. Indeed, another study has criticized the study by Zhang & Luck on exactly this point and did find an effect when using an experimental design with stronger incentives (Fougnie et al. 2016).

Another method to manipulate incentives is to vary the amount of potential reward across items within a display. For example, Klyszejko and colleagues performed a local change discrimination experiment in which the monetary reward for a correct response depended on which item was probed (Klyszejko et al. 2014). They found a positive relation between the amount of reward associated with an item and response accuracy, which indicates that subjects spent more resource on encoding items with larger potential reward. This incentive manipulation can be implemented by multiplying the behavioral cost function with an item-dependent factor *u_i_*, which modifies Eq. (11) to 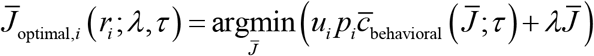. The coefficients *u_i_* and *p_i_* can be combined into a single “item relevance” coefficient *r_i_*=*u_i_p_i_*, and all theoretical results and predictions that we derived for *p_i_* now apply to *r_i_*.

A difference between the two discussed methods is that the former varied incentives within a trial and the latter across trials. However, both methods can be applied in both ways. A within-trial variant of the experiments by Zhang & Luck (2011) and Nassar et al. (2018) would be a *N*=2 task in which one of the items always has a low positive feedback threshold and the other a high one. Similarly, a between-trial variant of the experiment by Klyszejko et al. (2014) would be to scale the behavioral cost function of items with a factor that varies across trials or blocks, but is constant within a trial. Our model can be used to derive predictions for these task variants, which to our knowledge have not been reported on yet in the published literature.

### Neural mechanisms and timescale of optimization

Our results raise the question what neural mechanism could implement the optimal allocation policy that forms the core of our theory. Some form of divisive normalization (Bays 2014; Carandini & Heeger 2012) would be a likely candidate, which is already a key operation in neural models of attention (Reynolds & Heeger 2009) and visual working memory (Bays 2014; Wei et al. 2012). The essence of this mechanism is that it lowers the gain when set size is larger, without requiring explicit knowledge of the set size prior to the presentation of the stimuli. Consistent with the predictions of this theory, empirical work has found that the neural activity associated with the encoding of an item decreases with set size, as observed in for example the lateral intraparietal cortex (Churchland et al. 2008; Balan et al. 2008) and superior colliculus (Basso & Wurtz 1998). Moreover, the work by Bays (2014) has shown that a modified version of divisive normalization can account for the near-optimal distribution of resources across items with unequal probing probabilities. Since set size effects in our model are mediated by probing probability, its predicted set size effects can probably be accounted for by a similar mechanism.

Another question concerns the timescale at which the optimization takes place. In all experimental data that we considered here, the only factors that changed from trial to trial were set size (E1-E7) and probing probability (E7-E9). When we fitted the model, we assumed that the expected total cost in these experiments was minimized on a trial-by-trial basis: whenever set size or probing probability changed from one trial to the next, the computation of **J**_optimal_ followed this change. This assumption accounted well for the data and, as discussed above, previous work has shown that divisive normalization can accommodate trial-by-trial changes in set size and probing probability. However, can the same mechanism also accommodate changes in the optimal resource policy changes driven by other factors, such as the behavioral cost function, *c*_behavioral_(*ε*)? From a computational standpoint, divisive normalization is a mapping from an input vector of neural activities to an output vector, and the shape of this mapping depends on the parameters of the mechanism (such as gain, weighting factors, and a power on the input). Since the mapping is quite flexible, we expect that it can accommodate a near-optimal allocation policy for most experimental conditions. However, top-down control and some form of learning (e.g., reinforcement learning) are likely required to adjust the parameters of the normalization mechanism, which would prohibit instantaneous optimality after a change in the experimental conditions.

### Neural prediction

The total amount of resource that subjects spend on item encoding may vary non-monotonically with set size in our model. At the neural level, this translates to a prediction of a non-monotonic relation between population-level spiking activity and set size. We are not aware of any studies that have specifically addressed this prediction, but it can be tested using neuroimaging experiments similar to previously conducted experiments. For example, Balan et al. used single-neuron recording to estimate neural activity per item for set sizes 2, 4, and 6 in a visual search task (Balan et al. 2008). To test for the existence of the predicted non-monotonicity, the same recoding techniques can be used in a VWM task with a more fine-grained range of set sizes. Even though it is practically impossible to directly measure population-level activity, reasonable estimates may be obtained by multiplying single-neuron recordings with set size (under the assumption that an increase in resource translate to an increase in firing rate and not in an increase of neurons used to encode an item). A similar method can also assess the relation between an item’s probing probability and the spiking activity related to its neural encoding.

### Extensions to other domains

Our theory might apply beyond working memory tasks. In particular, it has been speculated that the selectivity of attention arises from a need to balance performance against the costs associated with spiking (Pestilli & Carrasco 2005; Lennie 2003). Our theory provides a normative formalism to test this speculation and may thus explain set size effects in attention tasks (Lindsay et al. 1968; Shaw 1980; Ma & Huang 2009).

Furthermore, developmental studies have found that that working memory capacity estimates change with age (Simmering & Perone 2012; Simmering 2012). Viewed from the perspective of our proposed theory, this raises the question why the optimal trade-off between behavioral and neural cost would change with age. A speculative answer is that a subject’s coding efficiency – formalized by the inverse of parameter *α* in Eq. (7) – may improve during childhood: an increase in coding efficiency reduces the neural cost per unit of precision, which shifts the optimal amount of resource to use for encoding to larger values. Neuroimaging studies might provide insight into whether and how coding efficiency changes with age, e.g. by estimating the amount of neural activity required per unit of precision in memory representations.

### Broader context

Our work fits into a broader tradition of normative theories in psychology and neuroscience (Table 2). The main motivation for such theories is to reach a deeper level of understanding by analyzing a system in the context of the ecological needs and constraints under which it evolved. Besides work on ideal-observer decision rules (Green & Swets 1966; Körding 2007; Geisler 2011; Shen & Ma 2016) and on resource-limited approximations to optimal inference (Gershman et al. 2015; Griffiths et al. 2015; Vul & Pashler 2014; Vul et al. 2009), normative approaches have also been used at the level of neural coding. For example, properties of receptive fields (Vincent et al. 2005; Liu et al. 2009; Olshausen & Field 1996), tuning curves (Attneave 1954; Barlow 1961; Ganguli & Simoncelli 2010), neural architecture (Cherniak 1994; Chklovskii et al. 2002), receptor performance (Laughlin 2001), and neural network modularity (Clune et al. 2013) have been explained as outcomes of optimization under either a cost or a hard constraint (on total neural firing, sparsity, or wiring length), and are thus mathematically closely related to the theory presented here. However, a difference concerns the timescale at which the optimization takes place: while optimization in the context of neural coding is typically thought to take place at the timescale over which the statistics of the environment change or a developmental timescale, the theory that we presented here could optimize on a trial-by-trial basis to follow changes in task properties.

**Table 2.**
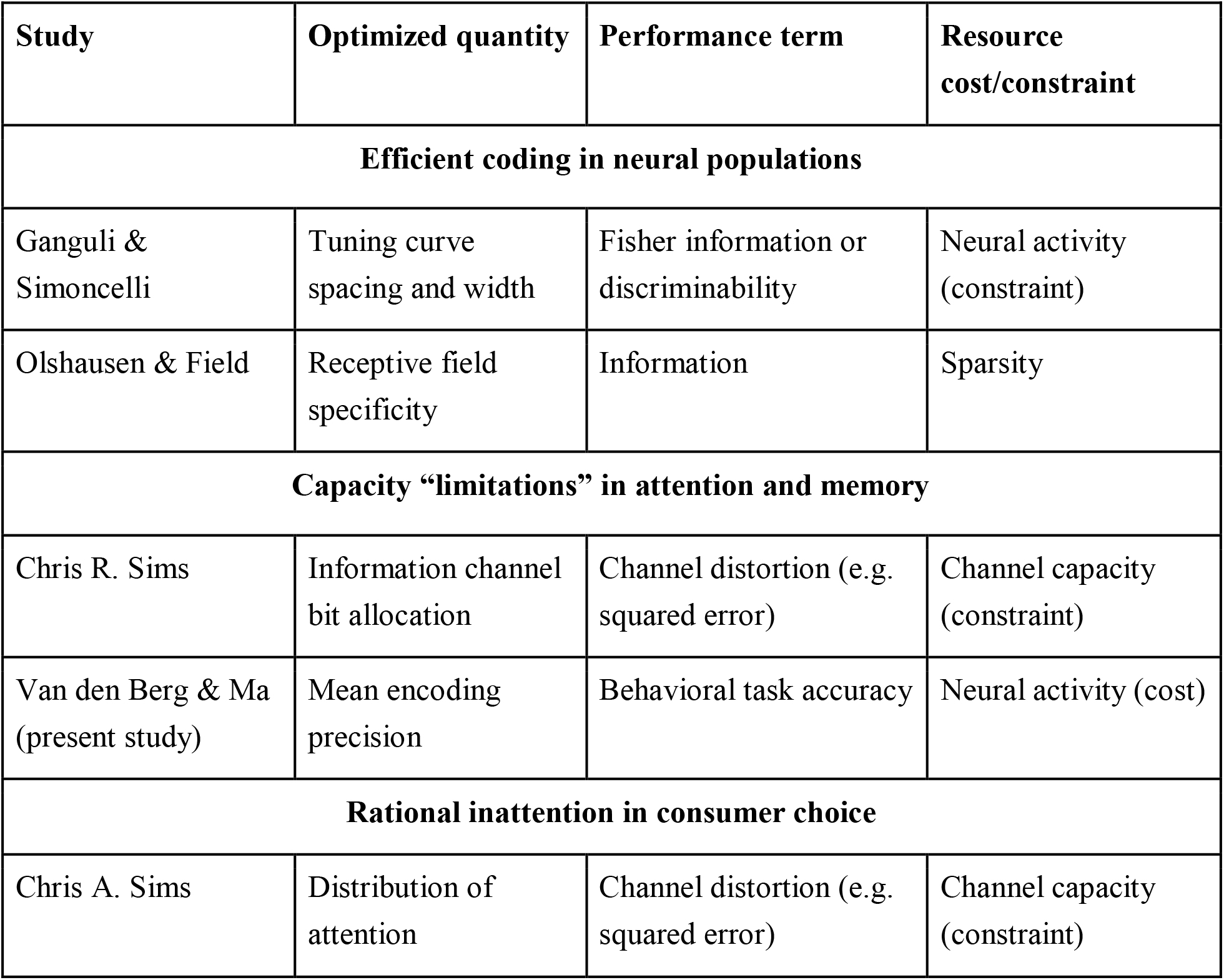
Examples of resource-rationality concepts in neuroscience, psychology, and economics.

| Study | Optimized quantity | Performance term | Resource cost/constraint |
| --- | --- | --- | --- |
| **Efficient coding in neural populations** |  |  |  |
| Ganguli & Simoncelli | Tuning curve spacing and width | Fisher information or discriminability | Neural activity (constraint) |
| Olshausen & Field | Receptive field specificity | Information | Sparsity |
| **Capacity “limitations” in attention and memory** |  |  |  |
| Chris R. Sims | Information channel bit allocation | Channel distortion (e.g. squared error) | Channel capacity (constraint) |
| Van den Berg & Ma (present study) | Mean encoding precision | Behavioral task accuracy | Neural activity (cost) |
| **Rational inattention in consumer choice** |  |  |  |
| Chris A. Sims | Distribution of attention | Channel distortion (e.g. squared error) | Channel capacity (constraint) |

We already mentioned the information-theory models of working memory developed by Chris R. Sims et al. A very similar framework has been proposed by Chris A. Sims in behavioral economics, who used information theory to formalize his hypothesis of “rational inattention”, i.e., the hypothesis that consumers make optimal decisions under a fixed budget of attentional resources that can be allocated to process economic data (Sims 2003). The model presented here differs from these two approaches in two important ways. First, similar to early models of visual working memory limitations, they postulate a fixed total amount of resources (formalized as channel capacity), which is a constraint rather than a cost. Second, even if it had been a cost, it would have been the expected value of a log probability ratio. Unlike neural spike count, a log probability ratio does not obviously map to a biologically meaningful cost on a single-trial level. Nevertheless, recent work has attempted to bridge rational inattention and attention in a psychophysical setting (Caplin et al. 2018).

## MATERIALS AND METHODS

### Data and code sharing

All data analyzed in this paper and model fitting code are available at [url to be inserted].

### Statistical analyses

Bayesian t-tests were performed using the JASP software package (JASP_Team 2017) with the scale parameter of the Cauchy prior set to its default value of 0.707.

### Model fitting

We used a Bayesian optimization method (Acerbi & Ma 2017) to find the parameter vector **θ**={*β*,*λ*,*τ*} that maximizes the log likelihood function, 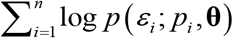, where *n* is the number of trials in the subject’s data set, *ε_i_* the estimation error on the *i*^th^ trial, and *p_i_* the probing probability of the probed item on that trial. To reduce the risk of converging into a local maximum, initial parameter estimates were chosen based on a coarse grid search over a large range of parameter values. The predicted estimation error distribution for a given parameter vector **θ** and probing probability *p_i_* was computed as follows. First, 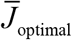 was computed by applying Matlab’s fminsearch function to Eq. (11). Thereafter, the gamma distribution over *J* (with mean 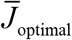 and shape parameter *τ*) was discretized into 20 equal-probability bins. The predicted (Von Mises) estimation error distribution was then computed under the central value of each bin. Finally, these 20 predicted distributions were averaged. We verified that increasing the number of bins used in the numerical approximation of the integral over J did not substantially affect the results.

### Model comparison using cross-validation

In the cross-validation analysis, we fitted the models in the same way as described above, but using only 80% of the data. We did this five times, each time leaving out a different subset of 20% the data (in the first run we left out trials 1, 6, 11, etc; in the second run we left out trials 2, 7, 12, etc; etc). At the end of each run, we used the maximum-likelihood parameter estimates to compute the log likelihood of the 20% of trials that were left out. These log likelihood values were then combined across the five runs to give an overall cross-validated log likelihood value for each model.

## ACKNOWLEDGMENTS

This work was funded by grant R01EY020958 from the National Institutes of Health, grant 2015-00371 by the Swedish Resarch Council, and grant INCA 600398 by Marie Sklodowska Curie Actions. We thank all authors of the papers listed in Table 1 for making their data available.

## Footnotes

1 The original benchmark set (van den Berg et al. 2014) contains 10 data sets. Three of those were published in papers that were later retracted and another one contains data at only two set sizes. While the model also accounts well for those data sets (Fig. S1 in Supplementary Information), we decided to exclude them from the main analyses.

2 In previous work (van den Berg et al. 2014) we have shown that the VP-A model outperforms basically any other descriptive VWM model, such as for example the slot-plus-averaging model (Zhang & Luck 2008). Hence, the finding that the resource-rational model is at equal footing with VP-A means that it outperforms most of the previously proposed descriptive models.

